# Extinction-resistant attention to long-term conditioned threat is indexed by selective visuocortical alpha suppression in humans

**DOI:** 10.1101/533141

**Authors:** Christian Panitz, Andreas Keil, Erik M. Mueller

## Abstract

While ERP studies have shown heightened early visual attention to conditioned threat, it is unknown whether this attentional prioritization is sustained throughout later processing stages and whether it is robust to extinction. To investigate sustained visual attention, we assessed visuocortical alpha suppression in response to conditioned and extinguished threat. We reanalysed data from *N* = 87 male participants that had shown successful long-term threat conditioning and extinction in self reports and physiological measures in a two-day conditioning paradigm. The current EEG time-frequency analyses on recall test data on Day 2 revealed that previously threat-conditioned vs. safety cues evoked stronger occipital alpha power suppression from 600 to 1200 ms. Notably, this suppression was resistant to previous extinction. The present study showed for the first time that threat conditioning enhances sustained modulation of visuocortical attention to threat in the long term. Long-term stability and extinction resistance of alpha suppression suggest a crucial role of visuocortical attention mechanisms in the maintenance of learned fears.

## Introduction

Learning to predict threat and safety based on environmental cues is fundamental for adaptive behaviour. A well-established paradigm for elucidating mechanisms of associative cue-outcome learning is classical threat conditioning (also fear/aversive conditioning) where conditioned stimuli are threat cues (CS+) when they signal an upcoming aversive event (unconditioned stimulus; US) and safety cues (CS−) when they signal the absence of an aversive event^1,2^. The acquired conditioned defensive response to the CS+ is expected to diminish again upon extinction training, i.e., repeated presentation of CS+ without US^3^.

Importantly, both conditioning and extinction memories need to be consolidated, retained, and recalled in future situations to allow adaptive responding in the long term. Successful short-term learning – i.e., acquisition and extinction of a conditioned response within an experimental session – is considered necessary but not sufficient for successful long-term learning. Such long-term learning effects are indicated by the successful recall of previously acquired and extinguished contingencies in a delayed test session^3^. Importantly, acquisition and extinction learning form separate memory traces and individuals may recall previously acquired threat memories without recalling extinction memories. In other words, they may show long-term extinction resistance, which is assumed to be key in the maintenance of fears^4^. In this case, despite repeated omission of aversive events, individuals keep showing robust conditioned threat responses over time.

Conditioned threat responses have different functions and manifest at various levels of central and autonomic physiology. Among these, increased attentive processing promotes capture of threat-relevant information, improving chances of successful defensive responding^5^. Supporting this notion, studies using visual evoked brain potentials in humans have found selectively heightened visual attention when viewing threat cues^6–11^. However, because visual evoked potentials to threat generally reflect early brain activity (i.e. < 500 ms)^12^, it is unknown, whether enhanced visual attention is sustained throughout later visual processing stages. Moreover, it is unclear, if heightened attention displays long-term resistance to extinction. These are clinically relevant questions given that enduring prioritization of threat processing may interfere with fear reduction through extinction and exposure therapy. The present study addressed these questions using a robust electrophysiological marker of sustained visual attention, suppression of visuocortical oscillatory activity in the alpha range.

Alpha oscillations (e.g., 8-12 Hz)^13,14^ indicate inhibition processes in neural populations^15,16^. Visually evoked posterior alpha suppression has been associated with increased excitability of early visual areas in the occipital cortex^17–20^ in response to increased attentional demands^21–24^. Moreover, posterior alpha power suppression has been found stronger in response to aversive vs. neutral/appetitive pictures^13,25,26^ (but also see ^27^), taken to reflect prioritized visual processing of threat information^5^.

In the present study, we reanalysed data from *N* = 87 participants that previously had shown successful long-term threat conditioning and extinction of CS ratings, evoked heart period and skin conductance responses assessed in a two-day differential threat conditioning paradigm with threat acquisition and extinction on one day and a critical recall test one day later^28^. For the current analyses, we estimated current source density of alpha-band activity at scalp sites consistent with visuocortical sources, to investigate (a) whether heightened visuocortical alpha suppression indexes selective visual attention to threat cues one day after conditioning, and (b) if conditioned alpha suppression is extinguished in the long term. For this purpose, we compared alpha power changes in response to previously extinguished vs. non-extinguished CS during the Day 2 recall test.

## Results

### Conditioning and extinction effects during Day 2 recall test

Means (and standard deviations) of alpha power percent change for the different CS types were as follows – CS+E: −4.47 (12.53), CS+N: −4.09 (12.95), CS-E: −1.17 (12.86), CS-N: −2.25 (14.84). The ANOVA showed a significant main effect of Contingency (*F*(1, 86) = 9.85, *p* = .002, η_p_^2^ = .103) as CS+ were followed by a stronger suppression of alpha power compared to CS− (also see Figure 1). Meanwhile, the main effect Extinction (*F*(1, 86) = 0.18, *p* = .669, η_p_^2^ = .002) and the Contingency x Extinction interaction (*F*(1, 86) = 1.29, *p* = .259, η_p_^2^ = .015) were not significant.

In line with the frequentist ANOVA, Bayesian ANOVA provided strongest evidence for a main effect of Contingency in the absence of other effects (BF_10_ = 26.0, all other models: BF_10_ < 3.3). In line with this pattern, Bayesian inclusion factors provided support for the inclusion of the main effect of Contingency (BF_Incl_ = 25.9) and against the inclusion of the main effect of Extinction (BF_Incl_ = 1 / 8.1) or the Contingency x Extinction interaction (BF_Incl_ = 1 / 4.0). Figure 2 shows time-frequency plots and topographic mapping of the Contingency effect on alpha power.

**Figure 1.**
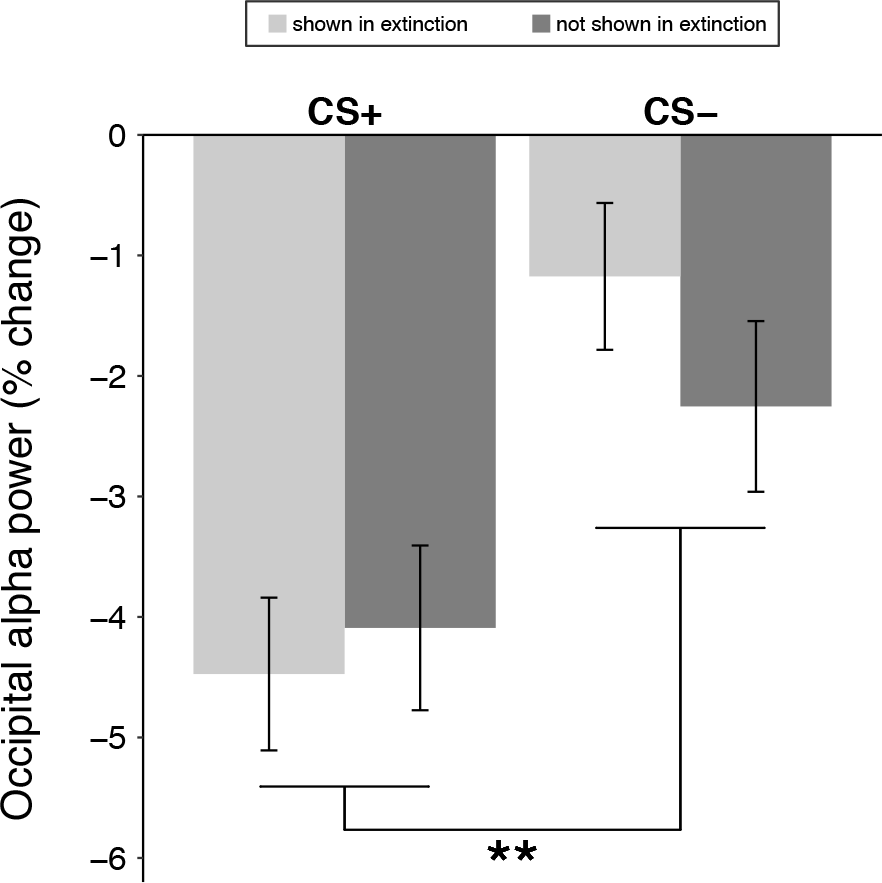
Conditioning effects on Day 2 alpha power. Mean alpha power (relative to baseline) at occipital sites in the time window of 500 – 1200 ms. Light bars represent CS presented during Day 1 extinction, dark bars represent CS not presented during Day 1 extinction. Error bars indicate SEM based on within-subject variance. **p < .01 for the main effect of Contingency.

**Figure 2.**
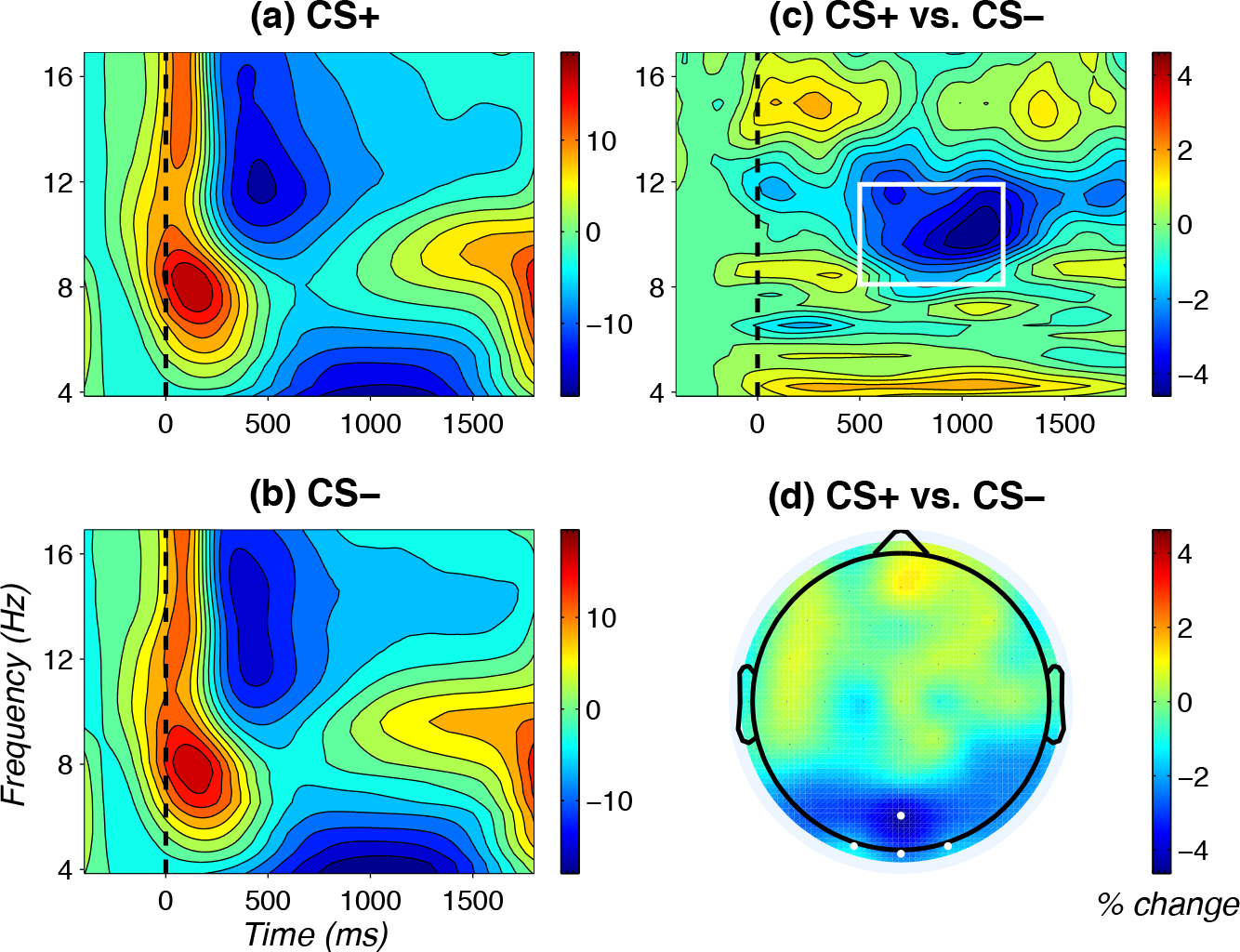
Main effect of Contingency on occipital alpha power. **(a)** and **(b)** Time-frequency plots for CS+ and CS−, respectively. Power values are % change relative to baseline (−400 to −200 ms) and averaged across Oz, POz, O1, and O2. **(c)** Time-frequency plot of the difference between CS+ and CS−, across Oz, POz, O1, and O2; the white rectangle indicates time (500 to 1200 ms) and frequency (8.1 to 11.9 Hz) windows for statistical analyses. **(d)** Topography of the difference between CS+ and CS− in the a priori defined time window (500 to 1200 ms). White dots depict the electrodes used for statistical analyses.

### Time course of Contingency effect

As suggested by permutation *t*-tests, viewing CS+ compared to CS− prompted significantly lower alpha power in the time window from 630 to 1230 ms post-CS (Figure 3), largely converging with our a priori window (500 to 1200 ms).

**Figure 3.**
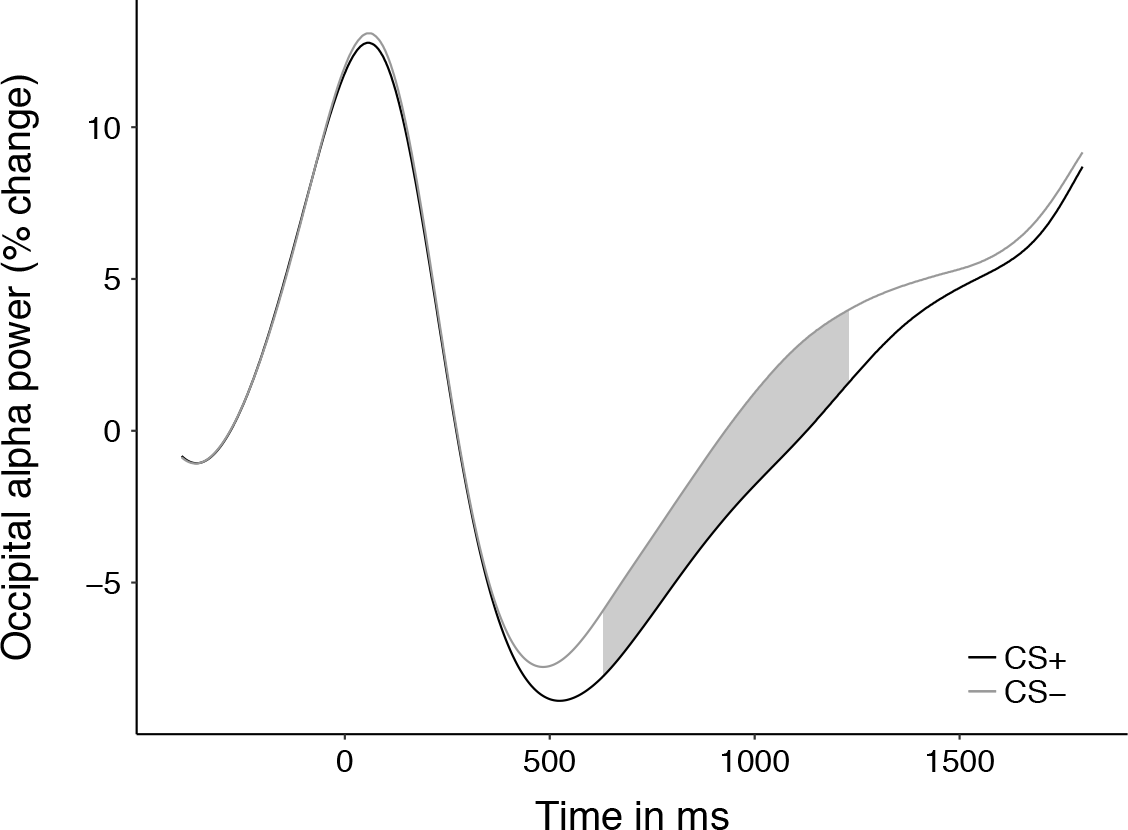
Time course of Contingency effect on mean occipital alpha power. Alpha power (relative to baseline) for collapsed CS+ and CS−, respectively, averaged across occipital sites. The grey-shaded area indicates significant differences between CS+ and CS− as determined by permutation testing (*p* < .05, two-sided).

## Discussion

In the present study, we investigated the role of heightened selective attention to threat cues in long-term threat conditioning and extinction. For this purpose, participants underwent a differential threat conditioning paradigm with acquisition and extinction on Day 1 and a test phase on Day 2 that allowed to assess long-term recall of extinguished and non-extinguished threat-conditioned responses. We found that, one day after threat conditioning, CS evoked a suppression of alpha power at occipital sites between 600 and 1200 ms, which was more pronounced for threat cues (i.e., CS+) compared to safety cues (i.e., CS−). Importantly, this conditioning effect was unaffected by Day 1 extinction training.

In the present sample, we observed robust alpha power suppression at occipital electrodes that was stronger for threat-conditioned CS+ than for CS−. This is the first time that occipital alpha power suppression is shown in response to conditioned threat cues. The present effects converge with previously shown alpha suppression to naturally threatening stimuli^13,25,26^ and can be interpreted as stronger disinhibition of early visuocortical areas facilitating attentional prioritization of behaviourally more relevant (i.e., threat) cues^22,24^. They are also consistent with previous ERP research showing threat-related prioritization of visuocortical processing^6–11,29,30^. Prioritization of threat is highly important for the successful choice of adaptive responding^5,31^, given limited capacities for information processing in the brain^32,33^.

Threat-potentiated alpha suppression was found in the time window from about 600 to 1200 ms post-CS. Therefore, this alpha suppression likely indicates *sustained* attention allocation to the threat cue via recruitment of visuocortical resources^13^. This process may be initiated after initial stimulus recognition and affective stimulus categorization as presumably indicated by early event-related potential^12,34^ and may reflect a more thorough analysis of threat cues. Taken together, the present results suggest occipital alpha as a promising new marker of sustained visuocortical attention modulation in response to conditioned threat cues.

Importantly, the stronger alpha suppression in response to CS+ was evident one day after initial learning, indicating successful long-term recall of selectively increased attention allocation to threat cues. To our knowledge, this is the first study to show such long-term threat conditioning effects on selective visual attention, stable across timely spaced situations. The long-term stability of the present alpha suppression suggests selective attention modulation in threat as a central feature of stable fear. While there is some evidence that selective attention to threatening information contributes to the long-term maintenance of fear^35,36^, the neural underpinnings of this link are not well understood. The present study suggests a mechanism in which differential activation of visuocortical areas is a temporally stable threat-conditioned response and underlies increased attentional processing of threat cues across time. Increased attentional threat processing may consolidate fearful behaviour via increased levels of state fear^36^ and by a reduced ability to disengage from threat cues at the cost of other potentially relevant stimuli^37–39^ – e.g., concurrent safety cues or cues for successful coping. Moreover, attentional biases may interact with negative expectancy, memory, and interpretation biases related to threat^40^. Future studies may use occipital alpha to investigate these and other potential mechanisms in order to elucidate attentional mechanisms of long-term fear maintenance.

In addition to long-term threat recall, we observed long-term extinction resistance of threat-conditioned alpha suppression. In other words, the initially threat-conditioned CS+ continued to evoke conditioned threat responses on Day 2, regardless of Day 1 fear extinction. The conclusion of extinction resistance was backed by Bayesian analyses, which favoured models excluding the extinction factor. Note that the present result pattern of extinction-resistant alpha suppression mirrors the pattern of long-term extinction-resistance of (a) rapid processing enhancements of threat-conditioned faces^30^ and (b) cardiac deceleration in the present sample^28^ as well as in previous studies^41,42^. Threat-evoked cardiac deceleration has been suggested to also reflect attentional processes, namely orienting in the face of imminent harm^43–45^. On the other hand, the present sample showed reduced differential skin conductance responses for extinguished (CS+E vs. CS-E) compared to non-extinguished (CS+N vs. CS-N) threat cues^28^, which indicates successful extinction recall of CS-US contingency awareness^46^. This suggests that attention-related processes in general, and occipital alpha suppression in particular, could be more extinction-resistant than other conditioned responses and may occur in a better-safe-than-sorry fashion. In other words, the cost of increased attentional engagement to invalid threat cues (i.e., false alarms) may be judged as significantly smaller than the cost for overlooking valid threat cues – even after repeated omission of harmful events.

As discussed earlier, alpha-related attentional processes may be crucial in the long-term maintenance of fears. The observed extinction resistance in the present study suggests that former threat cues continue drawing on attentional resources even in the absence of contingent reinforcement by harmful outcomes, i.e., even when cues are not followed by negative consequences anymore after initial learning. Moreover, occipital alpha suppression may prove useful for investigating the influence of threat-focused attention and attentional biases on the effectiveness of exposure therapy^47,48^ .

The limitations of the current study should be addressed. First, using the current paradigm, it cannot be ruled out that CS+ evoked stronger alpha suppression than CS− due to partial reinforcement and changing contingencies across phases, making predictions of US occurrence more difficult in CS+ vs. CS− and motivating participants to process CS+ more thoroughly. This, however, is unlikely to explain the present effects given that threat-depicting pictures, with no learning history, also evoked stronger alpha suppression in previous studies^13,25,26,49^. Second, we only used male participants in order to investigate conditioning and extinction without the influence of known sex differences^50^. As there also may be gender differences in visuocortical threat processing^51,52^, the present results should be replicated in females.

In the present study, we could show that conditioned visual threat cues evoke enhanced alpha suppression at occipital sites. Moreover, we showed for the first time that increased attention allocation to conditioned threat cues via sustained recruitment of early visuocortical areas is long-term stable and resistant to extinction. Future studies may use occipital alpha power to further elucidate mechanisms of visual attention in the development, maintenance, and extinction of fears.

## Methods

### Sample

We analysed data of a sample of *N* = 87 healthy, male participants (mean age: 23.7, SD: 3.85, range: 18-34), described in more detail elsewhere^28^. Participants were compensated 65 € (ca. 75 US$) for two experimental sessions (total 7 hours on two subsequent days). The study was conducted in accordance to the Declaration of Helsinki and was approved by the ethics committee of the German Psychological Association (DGPs). Informed consent was obtained from all participants at the beginning of the experiment.

### Experimental Design

#### Conditioning and Extinction paradigm

We employed a two-day differential threat conditioning and extinction paradigm^28,53^ (also see Figure 4). Briefly, on Day 1, participants underwent CS habituation, followed by an acquisition phase. Two CS (CS+E, CS+N) were paired with a US in 21 out of 45 trials (46.6 %), the other two CS (CS-E, CS-N) were never paired with the US (also 45 trials per CS). The extinction phase started exactly three hours after the end of acquisition and consisted of 40 CS+E and CS-E presentations (‘E’ standing for *presented during extinction phase)*. CS+N, CS-N, and US were not presented during the extinction phase (‘N’ standing for *not presented during extinction phase)*. Approximately 24 h after the extinction, participants returned for a Day 2 recall test which included 60 trials of each CS. No US were presented on Day 2.

#### Stimuli and trial structure

We used four different male faces with neutral expression from the Karolinska Directed Emotional Faces set^54^ as CS (pictures: AM10NES, AM13NES, AM31NES, BM08NES; also see Figure 4). Assignment of face stimuli to the different CS types was permutated and balanced across participants. The US was a 1 s white noise burst at 95 dB(A) delivered by a room speaker as we had previously shown that noise bursts are particularly well suited for threat conditioning with many trials^42^. In every trial, a fixation cross (1 s duration) was presented before participants saw the CS for 4 s. In paired trials the CS co-terminated with the US for 1 s. A black screen (jittered duration, 6-8 s) was presented between trials.

**Figure 4.**
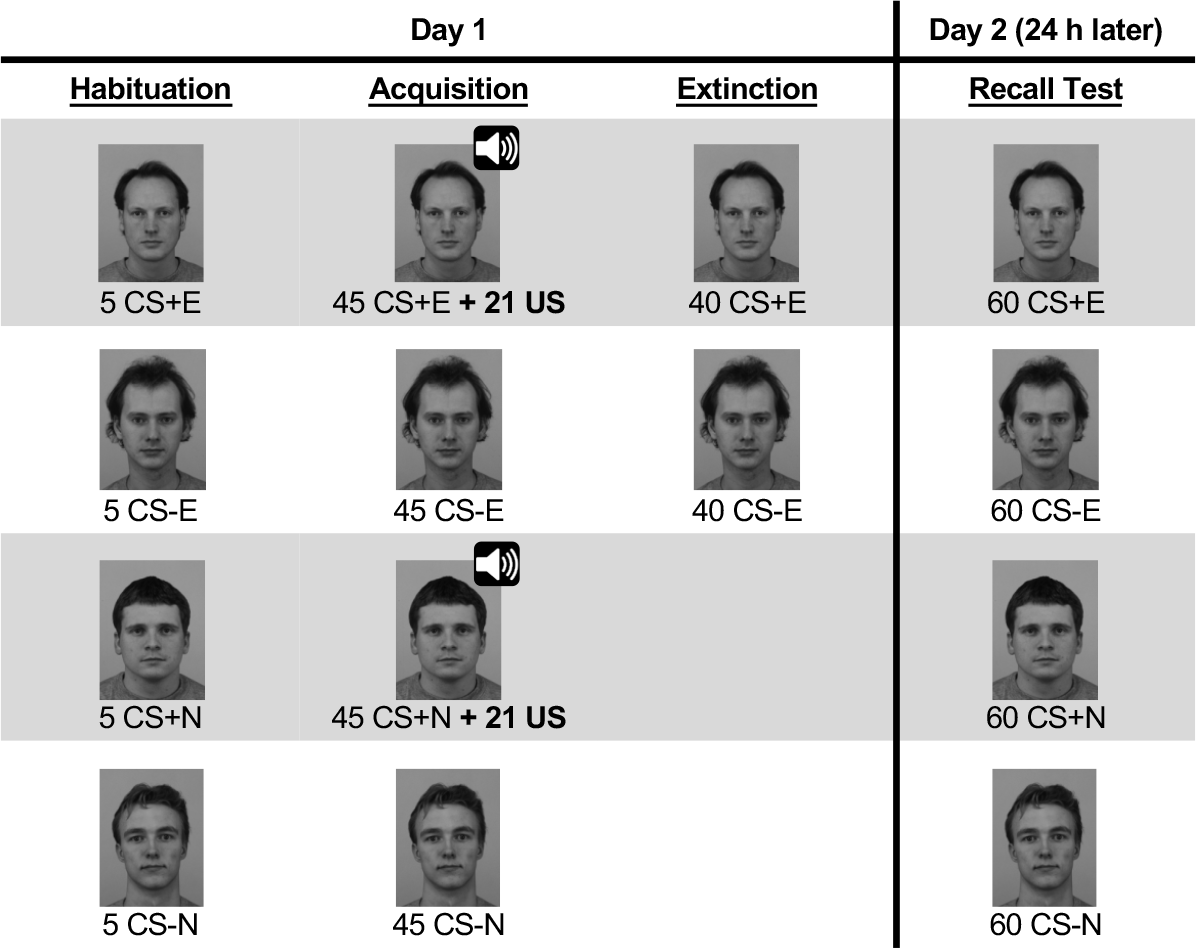
Conditioning and extinction paradigm. Face stimuli and number of presentations in the two-day differential threat conditioning and extinction paradigm. The US was only presented during acquisition, indicated by the speaker symbol. Assignment of different faces to CS type was permutated across participants. CS+E = extinguished CS+, CS+N = non-extinguished CS+, CS-E = CS− presented during extinction phase, CS-N = CS− not presented during extinction phase. Stimuli were presented in colour. KDEF stimuli IDs from top to bottom: AM10NES, AM13NES, AM31NES, BM08NES. Figure from ^28^.

### EEG recording and preprocessing

64-channel EEG was recorded with a QuickAmp72 amplifier and actiCAP active electrodes (Brain Products, Gilching, Germany) at 1000 Hz, with a 200 Hz online lowpass-filter and referenced against the average. EEG processing was performed in BrainVision Analyzer 2 (Brain Products, Gilching, Germany). The EEG was downsampled to 500 Hz, highpass-filtered (−3 dB at 0.1 Hz, 24 dB/oct., zero-phase IIR Butterworth filter) and notch-filtered (50 Hz, 5 Hz bandwidth, 16th order). Extended Infomax ICA^55^ was applied to the continuous data. Critical independent components reflective of horizontal and vertical eye movements, blinks, and cardiac artefacts were identified and removed by an experienced rater. To increase signal stationarity required for ICA, large EEG artefacts were removed manually, the signal was 0.5 Hz highpass-filtered for ICA only and the resulting weights were subsequently applied to the 0.1 Hz filtered data^56^. Segments with remaining artefacts were removed manually and channels with excessive amounts of bad data were interpolated (spline interpolation). Continuous EEG data were lowpass-filtered (30 Hz, filter specifications identical to highpass filter). After correcting marker latencies for monitor delay (33 ms), data were segmented relative to CS onset (−600 to 2000 ms).

To estimate current source density and improve the spatial specificity of the voltage map, the surface Laplacian was computed^57^ as implemented in BrainVision Analyzer 2 (spline order: 4, Legendre polynomial: 10, lambda: 1e-5). In order to facilitate comparison with previous studies, we also analysed the average-referenced scalp data. The results converged with the present results in the surface Laplacian data and are provided as supplementary material in the Open Science Framework (OSF) repository^58^.

### Wavelet analyses

Wavelet analyses were conducted in MATLAB 2013a (MathWorks, Natick, MA, USA) using established custom scripts^59^. First, EEG segments in the time domain were baseline-corrected (subtraction of the mean from −600 to −500 ms) and tapered with a cosine square window (20 samples rise/fall). Complex Morlet wavelets were applied with variable bandwidths (Morlet parameter = f/σ_f_ = 12) to compute power for frequency bands from 3.8 to 30.4 Hz in linear steps of 0.38 Hz. Power in each frequency band was baseline-corrected by dividing the signal by the mean amplitude between −400 ms and −200 ms relative to CS onset. Mean power of all discrete frequencies from 8.1 Hz to 11.9 Hz^13,14^ was used for statistical analyses on alpha.

### Statistical analyses

Statistical analyses were performed with R^60^ in the RStudio environment^61^. In order to assess the influence of Day 1 learning on occipital alpha during Day 2, we computed mean alpha power across Oz and adjacent electrodes (Oz, POz, O1, and O2) in the time window of 500 – 1200 ms^26,62^ after the CS. The resulting values were entered into an ANOVA with the within-subject factors Contingency (CS+ vs. CS−) and Extinction (extinguished vs. non-extinguished) using the *aov* function in R. Type I error level was set to α = .05. Distribution of mean alpha power across participants was sufficiently close to a normal distribution as indicated by low values of skewness and kurtosis for all CS types (|skewness| ≤ 0.46, |kurtosis| ≤ 1.12)^63^. In addition to null-hypothesis testing, we conducted Bayesian analyses as implemented in the *anovaBF* function in the BayesFactor package^64^ for R. We computed Bayes factors (100,000 iterations) for four different models (main effect Contingency only, main effect Extinction only, additive model of Contingency + Extinction main effects, complete model including both main effects and the interaction term^65^) compared to the null model (BF_10_). All models had equal prior probabilities and included a random effect term to account for between-subject variance. In a second step, we computed Bayesian inclusion factors (BF_Incl_) for each effect (i.e., Contingency, Extinction, and Contingency × Extinction) that indicate whether models including this effect are more likely to explain the data than matched models without this effect. More precisely, all models containing the effect of interest – but no higher-order interactions of this effect – were compared to matched models stripped of the effect (BAWS factor suggested by Sebastiaan Mathôt; also implemented in JASP 0.9^66^). As an example, for the Contingency effect, the models *Contingency only* and *Contingency + Extinction* were compared to the null model and the *Extinction only* model.

After using a predefined time window (500 to 1200 ms) for hypothesis testing, we conducted follow-up analyses on the exact time window of the Contingency effect based on the present data. We used permutation-controlled *t*-tests (adapted from the *t*_max_ approach from Blair and Karniski^67^), comparing alpha power (averaged across Oz, POz, O1, O2) at each time sample between both CS+ and both CS− (i.e., Contingency contrast). First, we randomly permutated the CS+ and CS− condition in each participant 1,000 times and computed *t*-values for each of the 1,300 time samples. Then, the tails of each permutation’s *t*-value distribution were determined and stored in a *t*_min_ and *t*_max_ distribution, respectively, each having 1,000 values corresponding to the 1,000 permutations. Finally, 2.5^th^ and 97.5^th^ percentiles from the resulting distribution of *t*_min_ and *t*_max_ values were used as critical *t*-values to compare empirical *t*-values against (i.e., α = .05, two-sided; *t*_crit_: CS+ < CS−: −2.59; CS+ > CS−: 2.64).

## Data availability

Data and R scripts for statistical analyses as well as preprocessed single-trial EEG data of the current study are available in the Open Science Framework (OSF) repository, osf.io/bfqjc.

## Acknowledgements

This study was supported by a grant of the German Research Foundation to Erik M. Mueller [DFG MU3535/2-1].

## Author Contributions

E.M.M. designed the research. C.P. collected the data. A.K. provided scripts for wavelet analyses. C.P. conducted wavelet and statistical analyses. All authors interpreted the results. C.P. drafted the manuscript and all authors edited the manuscript.

## Additional Information

The authors declare no competing interests.

## References

1. Lonsdorf, T. B. et al. Don’t fear ‘fear conditioning’: Methodological considerations for the design and analysis of studies on human fear acquisition, extinction, and return of fear. Neurosci. Biobehav. Rev. 77, 247–285 (2017).

2. Rescorla, R. A. & Wagner, A. R. A theory of Pavlovian conditioning: Variations in the effectiveness of reinforcement and nonreinforcement. in Classical conditioning II: Current research and theory (eds. Black, A. H. & Prokasy, W. F.) 64–99 (Appleton-Century-Crofts, 1972).

3. Myers, K. M. & Davis, M. Mechanisms of fear extinction. Mol. Psychiatry 12, 120–150 (2007).

4. Milad, M. R. & Quirk, G. J. Fear Extinction as a Model for Translational Neuroscience: Ten Years of Progress. Annu. Rev. Psychol. 63, 129–151 (2012).

5. Lang, P. J. & Bradley, M. M. Emotion and the motivational brain. Biol. Psychol. 84, 437–450 (2010).

6. Bublatzky, F. & Schupp, H. T. Pictures cueing threat: Brain dynamics in viewing explicitly instructed danger cues. Soc. Cogn. Affect. Neurosci. 7, 611–622 (2012).

7. Kastner-Dorn, A. K., Andreatta, M., Pauli, P. & Wieser, M. J. Hypervigilance during anxiety and selective attention during fear: Using steady-state visual evoked potentials (ssVEPs) to disentangle attention mechanisms during predictable and unpredictable threat. Cortex 106, 120–131 (2018).

8. Michalowski, J. M., Pané-Farré, C. A., Löw, A. & Hamm, A. O. Brain dynamics of visual attention during anticipation and encoding of threat- and safe-cues in spider-phobic individuals. 10, 1177–1186 (2015).

9. Wieser, M. J., Reicherts, P., Juravle, G. & von Leupoldt, A. Attention mechanisms during predictable and unpredictable threat — A steady-state visual evoked potential approach. Neuroimage 139, 167–175 (2016).

10. Gruss, L. F., Langaee, T. & Keil, A. The role of the COMT val158met polymorphism in mediating aversive learning in visual cortex. Neuroimage 125, 633–642 (2016).

11. Muench, H. M., Westermann, S., Pizzagalli, D. A., Hofmann, S. G. & Mueller, E. M. Self-relevant threat contexts enhance early processing of fear-conditioned faces. Biol. Psychol. 121, 194–202 (2016).

12. Miskovic, V. & Keil, A. Acquired fears reflected in cortical sensory processing: A review of electrophysiological studies of human classical conditioning. Psychophysiology 49, 1230–1241 (2012).

13. de Cesarei, A. & Codispoti, M. Affective modulation of the LPP and α-ERD during picture viewing. Psychophysiology 48, 1397–1404 (2011).

14. Bartsch, F., Hamuni, G., Miskovic, V., Lang, P. J. & Keil, A. Oscillatory brain activity in the alpha range is modulated by the content of word-prompted mental imagery. Psychophysiology 52, 727–735 (2015).

15. Klimesch, W., Sauseng, P. & Hanslmayr, S. EEG alpha oscillations: The inhibition-timing hypothesis. Brain Res. Rev. 53, 63–88 (2007).

16. Pfurtscheller, G. Functional brain imaging based on ERD / ERS. Vision Res. 41, 1257–1260 (2001).

17. Green, J. J. et al. Cortical and Subcortical Coordination of Visual Spatial Attention Revealed by Simultaneous EEG–fMRI Recording. J. Neurosci. 37, 7803–7810 (2017).

18. Scheeringa, R., Koopmans, P. J., van Mourik, T., Jensen, O. & Norris, D. G. The relationship between oscillatory EEG activity and the laminar-specific BOLD signal. Proc. Natl. Acad. Sci. 113, 6761–6766 (2016).

19. Iemi, L., Chaumon, M., Crouzet, S. M. & Busch, N. A. Spontaneous Neural Oscillations Bias Perception by Modulating Baseline Excitability. J. Neurosci. 37, 807–819 (2017).

20. Iemi, L. & Busch, N. A. Moment-to-Moment Fluctuations in Neuronal Excitability Bias Subjective Perception Rather than Strategic Decision-Making. eNeuro 5, 1–13 (2018).

21. Klimesch, W. Alpha-band oscillations, attention, and controlled access to stored information. Trends Cogn. Sci. 16, 606–617 (2012).

22. Foxe, J. J. & Snyder, A. C. The role of alpha-band brain oscillations as a sensory suppression mechanism during selective attention. Front. Psychol. 2, 154 (2011).

23. Ben-Simon, E. et al. The dark side of the alpha rhythm: FMRI evidence for induced alpha modulation during complete darkness. Eur. J. Neurosci. 37, 795–803 (2013).

24. Frey, J. N., Ruhnau, P. & Weisz, N. Not so different after all: The same oscillatory processes support different types of attention. Brain Res. 1626, 183–197 (2015).

25. Huster, R. J., Stevens, S., Gerlach, A. L. & Rist, F. A spectralanalytic approach to emotional responses evoked through picture presentation. Int. J. Psychophysiol. 72, 212–216 (2009).

26. Vagnoni, E., Lourenco, S. F. & Longo, M. R. Threat modulates neural responses to looming visual stimuli. Eur. J. Neurosci. 42, 2190–2202 (2015).

27. Uusberg, A., Uibo, H., Kreegipuu, K. & Allik, J. EEG alpha and cortical inhibition in affective attention. Int. J. Psychophysiol. 89, 26–36 (2013).

28. Panitz, C. et al. Fearfulness, neuroticism/anxiety, and COMT Val158Met in long-term fear conditioning and extinction. Neurobiol. Learn. Mem. 155, 7–20 (2018).

29. Thigpen, N. N., Bartsch, F. & Keil, A. The malleability of emotional perception: Short-term plasticity in retinotropic neurons accompanies the formation of perceptual biases to threat. J. Exp. Psychol. Gen. 146, 464–471 (2017).

30. Mueller, E. M. & Pizzagalli, D. A. One-year-old fear memories rapidly activate human fusiform gyrus. Soc. Cogn. Affect. Neurosci. 11, 308–316 (2016).

31. Bradley, M. M., Keil, A. & Lang, P. J. Orienting and emotional perception: facilitation, attenuation, and interference. Front. Psychol. 3, 493 (2012).

32. Chun, M. M., Golomb, J. D. & Turk-Browne, N. B. A Taxonomy of External and Internal Attention. Annu. Rev. Psychol. 62, 73–101 (2011).

33. Lavie, N. Distracted and confused?: Selective attention under load. Trends Cogn. Sci. 9, 75–82 (2005).

34. Schupp, H. T., Flaisch, T., Stockburger, J. & Junghöfer, M. Emotion and attention: event-related brain potential studies. Prog. Brain Res. 156, 31–51 (2006).

35. Richards, H. J., Benson, V., Donnelly, N. & Hadwin, J. A. Exploring the function of selective attention and hypervigilance for threat in anxiety. Clin. Psychol. Rev. 34, 1–13 (2014).

36. Van Bockstaele, B. et al. A Review of Current Evidence for the Causal Impact of Attentional Bias on Fear and Anxiety. Psychol. Bull. 140, 682–721 (2014).

37. Fox, E. et al. Attentional bias for threat: Evidence for delayed disengagement from emotional faces. Cogn. Emot. 16, 355–379 (2002).

38. Georgiou, G. A. et al. Focusing on fear: Attentional disengagement from emotional faces. Vis. cogn. 12, 145–158 (2005).

39. Rinck, M., Reinecke, A., Ellwart, T., Heuer, K. & Becker, E. S. Speeded Detection and Increased Distraction in Fear of Spiders: Evidence From Eye Movements. J. Abnorm. Psychol. 114, 235–248 (2005).

40. Aue, T. & Okon-Singer, H. Expectancy biases in fear and anxiety and their link to biases in attention. Clin. Psychol. Rev. 42, 83–95 (2015).

41. Panitz, C., Hermann, C. & Mueller, E. M. Conditioned and extinguished fear modulate functional corticocardiac coupling in humans. Psychophysiology 52, 1351–1360 (2015).

42. Sperl, M. F., Panitz, C., Hermann, C. & Mueller, E. M. A pragmatic comparison of noise burst and electric shock unconditioned stimuli for fear conditioning research with many trials. Psychophysiology 53, (2016).

43. Löw, A., Weymar, M. & Hamm, A. O. When Threat Is Near, Get Out of Here: Dynamics of Defensive Behavior During Freezing and Active Avoidance. Psychol. Sci. 26, 1706–1716 (2015).

44. Wendt, J., Löw, A., Weymar, M., Lotze, M. & Hamm, A. O. Active avoidance and attentive freezing in the face of approaching threat. Neuroimage 158, 196–204 (2017).

45. Roelofs, K. Freeze for action: Neurobiological mechanisms in animal and human freezing. Philos. Trans. R. Soc. B 372, 20160206 (2017).

46. Hamm, A. O. & Vaitl, D. Affective learning: Awareness and aversion. Psychophysiology 33, 698–710 (1996).

47. Barry, T. J., Vervliet, B. & Hermans, D. An integrative review of attention biases and their contribution to treatment for anxiety disorders. Front. Psychol. 6, 968 (2015).

48. Mogg, K. & Bradley, B. P. Anxiety and Threat-Related Attention: Cognitive-Motivational Framework and Treatment. Trends Cogn. Sci. 22, 225–240 (2018).

49. Güntekin, B. & Basar, E. Emotional face expressions are differentiated with brain oscillations. Int. J. Psychophysiol. 64, 91–100 (2007).

50. Merz, C. J. et al. Neuronal correlates of extinction learning are modulated by sex hormones. Soc. Cogn. Affect. Neurosci. 7, 819–830 (2012).

51. Pintzinger, N. M., Pfabigan, D. M., Pfau, L., Kryspin-Exner, I. & Lamm, C. Temperament differentially influences early information processing in men and women: Preliminary electrophysiological evidence of attentional biases in healthy individuals. Biol. Psychol. 122, 69–79 (2017).

52. Sass, S. M. et al. Time course of attentional bias in anxiety: Emotion and gender specificity. Psychophysiology 47, 247–259 (2010).

53. Mueller, E. M., Panitz, C., Hermann, C. & Pizzagalli, D. A. Prefrontal Oscillations during Recall of Conditioned and Extinguished Fear in Humans. J. Neurosci. 34, 7059–7066 (2014).

54. Lundqvist, D., Flykt, A. & Öhman, A. The Karolinska Directed Emotional Faces - KDEF. CD ROM from the Department of Clinical Neuroscience, Psychology Section, Karolinska Institutet. (1998).

55. Lee, T. W., Girolami, M. & Sejnowski, T. J. Independent component analysis using an extended infomax algorithm for mixed subgaussian and supergaussian sources. Neural Comput. 11, 417–441 (1999).

56. Winkler, I., Debener, S., Muller, K.-R. & Tangermann, M. On the influence of high-pass filtering on ICA-based artifact reduction in EEG-ERP. in 2015 37th Annual International Conference of the IEEE Engineering in Medicine and Biology Society (EMBC) 4101–4105 (2015). doi:10.1109/EMBC.2015.7319296

57. Perrin, F., Pernier, J., Bertrand, O. & Echallier, J. F. Spherical splines for scalp potential and current density mapping. Electroencephalogr. Clin. Neurophysiol. 72, 184–187 (1989).

58. Panitz, C., Keil, A. & Mueller, E. M. Extinction-resistant attention to long-term conditioned threat is indexed by selective visuocortical alpha suppression - Open Data and Materials. (Available at: osf.io/bfqjc, 2019).

59. Keil, A., Stolarova, M., Moratti, S. & Ray, W. J. Adaptation in human visual cortex as a mechanism for rapid discrimination of aversive stimuli. Neuroimage 36, 472–479 (2007).

60. R Core Team. R: A Language and Environment for Statistical Computing. (R Foundation for Statistical Computing, 2018).

61. RStudio Team. RStudio: Integrated Development Environment for R. (RStudio, Inc., 2016).

62. Chien, J. H. et al. Oscillatory EEG Activity Induced by Conditioning Stimuli During Fear Conditioning Reflects Salience and Valence of these Stimuli More than Expectancy. Neuroscience 346, 81–93 (2018).

63. Curran, P. J., West, S. G. & Finch, J. F. The Robustness of Test Statistics to Nonnormality and Specification Error in Confirmatory Factor Analysis. Psychol. Methods 1, 16–29 (1996).

64. Morey, R. D. & Rouder, J. N. BayesFactor: Computation of Bayes Factors for Common Designs. (2018).

65. Rouder, J. N., Engelhardt, C. R., McCabe, S. & Morey, R. D. Model comparison in ANOVA. Psychon. Bull. Rev. 23, 1779–1786 (2016).

66. JASP Team. JASP (Version 0.9)[Computer Software]. (2018).

67. Blair, R. C. & Karniski, W. An alternative method for significance testing of waveform difference potentials. Psychophysiology 30, 518–524 (1993).

